# History matters: thermal environment before, but not during wasp attack determines the efficiency of symbiont-mediated protection

**DOI:** 10.1101/2022.09.30.510345

**Authors:** Jordan E. Jones, Gregory D. D. Hurst

## Abstract

The outcome of natural enemy attack in insects is commonly impacted by the presence of defensive microbial symbionts residing within the host. Beyond their presence, the outcome of the interaction can also depend on genetic and environmental factors. The thermal environment is a key factor known to affect symbiont-mediated traits in insects, including their ability to defend against natural enemy attack. Cooler temperatures, for instance, have been previously shown to reduce *Spiroplasma*-mediated protection in *Drosophila*. Here, we dissect the effect of the thermal environment on *Spiroplasma*-mediated protection against *Leptopilina boulardi* in *Drosophila melanogaster* by examining the effect of temperature before, during and after wasp attack on fly survival and wasp success. We observed that the developmental temperature of the mothers’ of attacked larvae, and not the temperature of the attacked larvae themselves during or after wasp attack, strongly determines the protective influence of *Spiroplasma. Spiroplasma*-mediated fly survival was found to be weaker when parental flies were reared at 21°C before their larvae were exposed to wasps compared to larvae derived from parental flies reared at 23°C or 25°C. Contrastingly, there was no effect of thermal environment on protection when mothers were reared at 25°C, and their progeny exposed to lower temperatures during and after wasp attack. The effect of developmental temperature on *Spiroplasma*-mediated protection is likely mediated by reduced *Spiroplasma* titre combined, at cooler temperatures, with segregation of infection. These results indicate the historical thermal environment is a stronger determinant of protection than current environment, and that protective capacity is partly an epigenetic trait.

## Introduction

Heritable microbial symbionts are now widely recognised to be key players in the outcome of insect-parasite interactions, providing an important line of defence against attack. The diversity of defensive symbiont associations described has increased dramatically over recent years with symbionts affording protection against a wide range of natural enemies including parasitoid wasps (Oliver *et al*., 2003; Xie *et al*., 2010), fungal pathogens (Scarborough *et al*., 2005; Lukasik *et al*., 2013), nematodes (Jaenike *et al*., 2010) and ssRNA viruses (Hedges *et al*., 2008; Teixeira *et al*., 2008). Microbial symbionts and their defensive properties are also important in terms of their application. For instance, the antiviral properties of *Wolbachia* have been utilised in natural populations of mosquitoes to protect humans from diseases such as dengue and Zika (O’Neill *et al*., 2018; Utarini *et al*., 2021). Protective traits may also be important in biological control programmes (Rossbacher and Vorburger, 2020). Thus, it is important to gain a general understanding of what factors can impact the protection afforded by defensive bacterial symbionts.

Beyond the mere presence or absence of the defensive symbiont, the outcome of natural enemy attack can depend on the strain of all players in the interaction (Xie *et al*., 2010; Schmid *et al*., 2012; Chrostek *et al*., 2013; Cayetano and Vorburger, 2013). However, in comparison to host, parasite and symbiont genetics, much less is known about how environmental factors contribute to the penetrance of symbiont-mediated defence. A few studies have observed thermal sensitivity of symbiont-mediated protection. In the pea aphid, *H. defensa*-mediated protection against *A. ervi* has been repeatedly shown to be negatively impacted, or fail, at warmer temperatures relative to cooler controls, with even a modest temperature rise of 2.5°C enough to reduce the strength of *H. defensa*-mediated protection (Bensadia *et al*., 2006; Doremus *et al*., 2018; Higashi *et al*., 2020). In contrast to the studies above, *Spiroplasma*-mediated protection against *Leptopilina heterotoma* in *Drosophila hydei* completely failed at a cooler temperature of 18°C compared to 25°C (Corbin *et al*., 2021).

A key issue with respect to thermal impacts on protection strength is defining the sensitive period in which temperature impacts the phenotype. Whilst intuition might indicate the temperature at the point of attack is of paramount importance, this is not necessarily the case. For instance, in *D. melanogaster, Wolbachia* conferred strong protection against *Drosophila* C virus when flies were reared from egg to adult through 25°C pre-infection, while this protective effect was lost when flies were reared through a cooler temperature of 18°C (Chrostek *et al*., 2021). Here, there was a clear effect of historical thermal environment on protection, mirroring epigenetic influences on phenotype observed in other host-symbiont interactions (e.g. Dyer and Jaenike, 2005; Layton *et al*., 2019).

In this study, we dissect the thermal sensitivity of *Spiroplasma*-mediated protection in *D. melanogaster* in terms of sensitivity to cool environments at different timepoints. *Spiroplasma* is a facultative symbiont of *Drosophila* and has been shown to protect against parasitoid wasps and nematodes (Xie *et al*., 2010; Jaenike *et al*., 2010), and the strain in *D. melanogaster* combines protection phenotype with male-killing. Previous studies on this system have concluded that the strain of wasp and *Spiroplasma* are all important components for the outcome of *Spiroplasma*-mediated protection (Jones and Hurst, 2020a; Jones and Hurst, 2020b). Whilst influence of temperature on the protective phenotype have not been established, vertical transmission of *Spiroplasma* in *D. melanogaster* fails at 16.5°C, indicating cool sensitivity of the symbiosis (Montenegro and Klaczko, 2004).

Within our analysis, we consider the importance of thermal environment at two stages: the pre-attack stage (exposure of the parental generation, eggs and L1 larvae before attack) and the attack and protection stage (attack of L1 /L2 larvae and their subsequent development into adults). From this analysis, we can determine the degree to which it is the thermal environment during parasitism which is important, compared to historical impacts arising before parasitism. We also examined how these thermal regimes impact *Spiroplasma* titre, an aspect of symbiosis which can potentially underpin the strength of both protective and male-killing phenotypes.

## Materials and methods

### Insect strains and maintenance

Experiments used the MSRO-Ug strain *Spiroplasma* strain originally collected from Namulonge, Uganda in 2005 (Pool *et al*., 2006); this strain was transinfected onto a Canton-S background in 2020 (Jones and Hurst, 2020b). MSRO-Ug Canton-S stocks and Canton-S stocks lacking *Spiroplasma* were maintained on corn meal agar (10 g agarose, 85 g sugar, 60 g maize meal, 40 g autolysed yeast in a total volume of 1 L, to which 25 mL 10% Nipagin dissolved in 100% ethanol was added) at 25°C on a 12:12 light:dark cycle.

Parasitism assays utilised the inbred Lb17 strain of *Leptopilina boulardi*, initially collected in Winters, California in 2002 (Schlenke *et al*., 2007). Wasp stocks were maintained on second instar Canton-S larvae at 25°C on a 12:12 light:dark cycle. After emergence, wasps were maintained in vials containing sugar yeast medium (20 g agarose, 100 g sugar, 100 g autolysed yeast in a total volume of 1 L, to which 30 mL 10% Nipagin dissolved in 100% ethanol and 3 mL propionic acid was added) supplemented with a Flugs© (Flystuff, Genesee Scientific) moistened with honey water and allowed to mature and mate for at least 7 days prior to exposure to *D. melanogaster* L2 larvae.

### The effect of temperature on *Spiroplasma*-mediated fly survival and wasp mortality

We dissected the effect of thermal environment at different timepoints on *Spiroplasma* mediated protection. Wasp attack assays were performed across three different thermal environments (21°C, 23°C or 25°C) over two different temporal periods (‘pre-attack’ and ‘attack and protection’) (Figure 1). The ‘pre-attack’ regime represented the effect of temperature (21°C, 23°C or 25°C) on maternal egg to adult development on *Spiroplasma*-mediated fly survival of their wasp exposed offspring (this also included 3-5 days of early adulthood and one day of egg laying). The ‘attack and protection’ temperature regime determined the effect of temperature (21°C, 23°C or 25°C) on *Spiroplasma*-mediated fly survival at the point of wasp attack on L2 larvae and the subsequent development following wasp attack during which *Spiroplasma* is actively defending against wasp larvae development. Finally, the effect of a constant experimental temperature across the ‘pre-attack’ stage and the ‘attack and protection’ stage of either 21°C, 23°C or 25°C was also completed.

**Figure 1:**
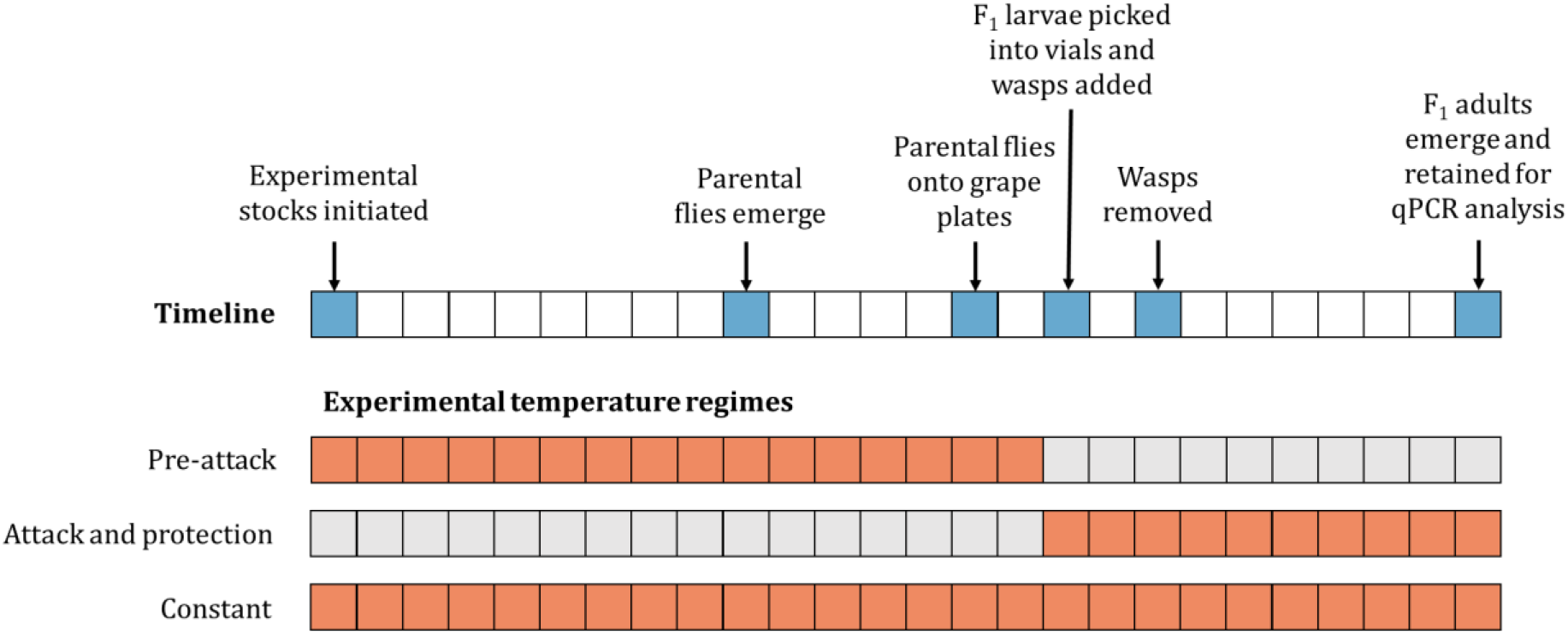
Experimental design showing the wasp attack assay timeline and the three experimental temperature regimes conducted. For each regime, orange squares indicate when flies were subject to an experimental temperature of either 21°C, 23°C or 25°C. Grey squares indicate when all flies were subject to a standard 25°C temperature. Timeline squares represent a single day based on a fly generation at 25°C.

To this end, MSRO-Ug stocks and uninfected isogenic control Canton-S stocks were initiated by placing 3 females and 2 males into ASG vials supplemented with yeast granules and placed at their experimental temperatures to develop (MSRO-Ug females were given 2 Canton-S males to mate). Adults were removed from vials 4 days after egg laying. As the development time of *Drosophila melanogaster* increases at lower temperatures (21°C = 15 days; 23°C = 12 days; 25°C = 10 days) the start of the pre-attack and constant temperature regime was staggered so that parasitisation was conducted at the same time to control for variation in wasp oviposition. All founding MSRO-Ug females were 6 days old at the point of egg laying to control for any age-mediated effects of *Spiroplasma* titre.

Adult progeny from founder females were allowed to emerge over 3 days before being collected into vials containing sugar yeast medium and supplemented with live yeast paste. MSRO-Ug female flies were given Canton-S males to mate. After 2 days of mating, flies were transferred into cages and allowed to continue to mate and lay eggs on a grape Petri dish painted with live yeast paste for 24 h at their experimental temperature. 24 hours later, first instar larvae were picked from the grape plates into the experimental vials at 30 larvae per vial. Twelve treatments were formed in total for each temperature regime with 10 replicate vials per treatment [3 temperatures (21°C, 23°C or 25°C) x 2 symbiont treatments (infected/uninfected) x 2 wasp treatments (wasp/no wasp)]. Five female wasps were transferred into the wasp treatment vials and allowed 48 h to attack before being removed. All vials were maintained at their experimental temperature on a 12:12 light:dark cycle. For each vial, the number of pupae, emerging wasps and emerging flies was counted. The sex of each emerging fly was also recorded to determine temperature effects on the male-killing phenotype and to indicate loss of *Spiroplasma* infection. *Spiroplasma*-infected wasp-attacked and no wasp control flies emerging from the constant temperature regime were collected at 1 day old and frozen at −20°C for subsequent qPCR analysis.

### The effect of temperature and wasp attack on *Spiroplasma* titre

The effect of temperature and wasp attack on *Spiroplasma* titre was also assessed. DNA template was prepared from individual flies emerging from the experiment and *Spiroplasma* titre was estimated by quantitative PCR (qPCR). To this end, DNA was extracted from 1 day old individual female flies (15-20 flies per treatment) using the Phenol-Chloroform method. DNA concentrations and quality were determined using NanoDrop ND-1000 spectrophotometer. All samples were stored at −20°C. Real-time qPCRs were carried out for the *dnaA* gene (*Spiroplasma*) using the primers, DnaA109F (5’-TTA AGA GCA GTT TCA AAA TCG GG-3’), and DnaA246R (5’-TGA AAA AAA CAA ACA AAT TGT TAT TAC TTC-3’) and for the *RPS17* gene (*Drosophila melanogaster* reference) using the primers Dmel.rps17F (5’-CAC TCC CAG GTG CGT GGT AT-3’) and Dmel.rps17R (5’-GGA GAC GGC CGG GAC GTA GT-3’), using the LightCycler 480 (Roche) (Anbutsu and Fukatsu, 2003; Osborne *et al*., 2009). Each reaction consisted of 6 μl of PowerUp™ SYBR™ Green Master Mix (ThermoFisher), 0.5 μl of each primer solution at 3.6 μm and 5 μl of diluted DNA, with three technical replicates per reaction. Average Ct values were calculated using the average across 2 or 3 replicates that were within a standard deviation of 0.5. Relative amounts of *Spiroplasma* were calculated using the Pfaffl Method (Pfaffl, 2001).

### The effect of temperature on wasp oviposition

Differences in fly survival in the face of wasp attack may be a consequence of changes in wasp oviposition behaviour at differing temperatures. To determine whether wasp oviposition differed with environmental temperature, we compared the number of wasps (eggs and larvae) per fly larva after a 48 hour exposure period to Lb17 wasps at 21°C, 23°C and 25°C. To this end, the same protocol was followed to obtain fly larvae as the wasp attack assay described above. Five female wasps were placed into each vial containing 30 L2 *D. melanogaster* Canton-S larvae for 48 hours. Immediately after wasp removal, approximately 5 fly larvae from each of the 6 replicate vials were dissected under a microscope and the number of wasp eggs and/or larvae present were counted.

### Statistical analysis

All statistical analyses were done in *RStudio* (RStudio Team, 2020) using the statistical software, *R* version 4.0.2 (R Core Team, 2022). All figures were produced using *ggplot2* (Wickham, 2016).

Fly and wasp survival data were analysed by fitting a generalised linear model with binomial or quasibinomial errors to account for overdispersion where necessary. A two-way factorial analysis of deviance was performed to test the effect of temperature (21°C, 23°C and 25°C) and *Spiroplasma* infection as well as the two-way interaction on fly and wasp survival. The *‘Anova’* function in the *‘car’* package v.3.1-0 with *F* tests were used (Fox and Weisberg, 2019).

*Spiroplasma* titre data were analysed by fitting a generalised linear model with gamma distribution. A two-way factorial analysis of deviance was performed to test the effect of temperature (21°C, 23°C and 25°C) and wasp attack as well as the two-way interaction on relative *Spiroplasma* titre of flies. The *‘Anova’* function in the *‘car’* package v.3.1-0 with Wald chi-squared tests were used (Fox and Weisberg, 2019).

Wasp oviposition data was tested for normality and subsequently analysed using the Kruskal-Wallis rank sum test.

Sample sizes for all experiments are found in Table S1.

## Results

### Interaction between thermal environment, Spiroplasma infection and fly survival in the absence of wasp attack

In the absence of wasps, temperature had no significant effect on the fly larva-to-adult survival across temperature regimes (Table 1). The mean larva-to-adult fly survival was > 68% across all treatments (Figure 2A; B; C). *Spiroplasma* infection had a significant effect on the fly larva-to-adult survival across all temperature regimes in the absence of wasps (Table 1). The mean fly larva-to-adult survival of *Spiroplasma*-infected flies was lower than *Spiroplasma*-uninfected flies in the pre-attack and constant temperature regime (pre-attack: S+ mean fly survival = 73.4%, n = 30 vials vs. S-mean fly survival = 84.4%, n = 28; constant: S+ mean fly survival = 76.8%, n = 30 vs. S-mean fly survival = 84.5%, n = 28). However, in the constant temperature regime, *Spiroplasma*-infected fly survival was higher at 21°C resulting in a significant interaction between *Spiroplasma* infection and temperature (21°C constant: S+ mean fly survival = 82.3%, n = 10 vials vs. S-mean fly survival = 80.7%, n = 10; Table 1). In contrast to the pre-attack and constant temperature regimes, the mean fly larva-to-adult survival of *Spiroplasma*-infected flies was higher than *Spiroplasma*-uninfected flies in the attack and protection temperature regime (S+ mean fly survival = 88.2%, n = 24 vs. S-mean fly survival = 79.9%, n = 26).

**Table 1:** Analysis of deviance table for fly larva-to-adult survival for the ‘pre-attack’, ‘attack and protection’ and ‘constant’ temperature regimes in the no wasp control treatment.

| **Pre-attack** | Sum Sq | df | *F* | *p*-value |
| --- | --- | --- | --- | --- |
| Temperature | 9.50 | 2 | 2.58 | 0.085 |
| *Spiroplasma* infection | 31.80 | 1 | 17.29 | <0.001 |
| Temperature * *Spiroplasma* infection | 1.24 | 2 | 0.34 | 0.714 |
| Residuals | 95.64 | 52 |  |  |
| **Attack and protection** |  |  |  |  |
| Temperature | 12.66 | 2 | 2.24 | 0.118 |
| *Spiroplasma* infection | 20.42 | 1 | 7.23 | 0.010 |
| Temperature * *Spiroplasma* infection | 11.37 | 2 | 2.01 | 0.146 |
| Residuals | 124.21 | 44 |  |  |
| **Constant** |  |  |  |  |
| Temperature | 1.23 | 2 | 0.32 | 0.728 |
| *Spiroplasma* infection | 16.64 | 1 | 8.68 | 0.005 |
| Temperature * *Spiroplasma* infection | 19.33 | 2 | 5.04 | 0.010 |
| Residuals | 99.67 | 52 |  |  |

*Note: A generalised linear model with logit link and quasibinomial fit was applied. (Pre-attack): The dispersion parameter is 1.839. (Attack and protection): The dispersion parameter is 2.823. (Constant): The dispersion parameter is 1.917.*

**Figure 2:**
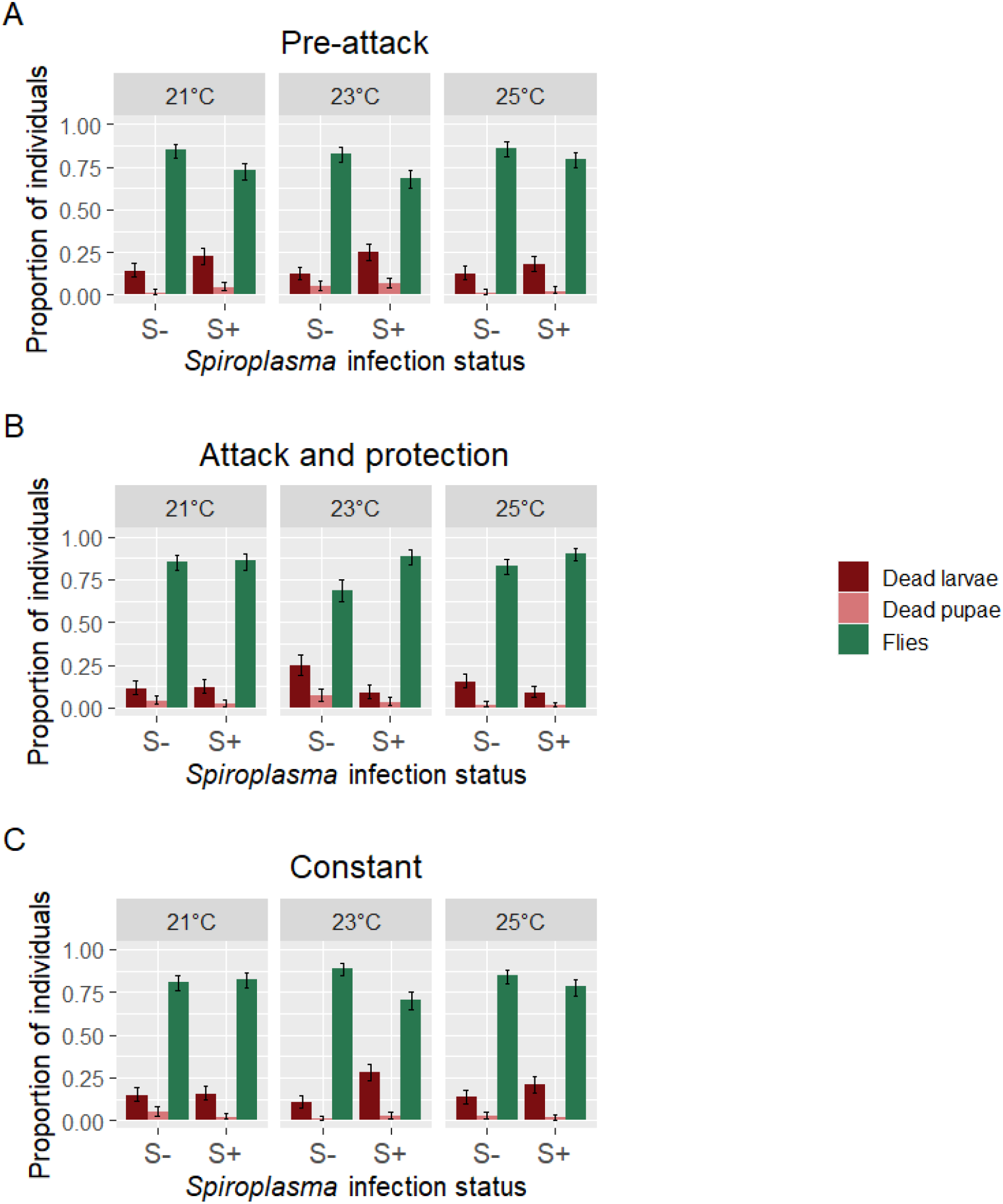
Proportion of dead larvae (red), dead pupae (pink) and emerging flies (green) for *Spiroplasma*-uninfected (S-) and -infected (S+) F1 *Drosophila melanogaster* under the no wasp control ‘pre-attack’ (A), ‘attack and protection’ (B) and ‘constant’ (C) temperature regimes. Error bars represent 95% binomial confidence intervals.

### Interaction between thermal environment, Spiroplasma infection and fly survival in the presence of wasp attack

In the presence of wasps, temperature was found to have a significant effect on the fly larva-to-adult survival, in both the ‘pre-attack’ and the ‘constant’ temperature regimes (Table 2). As very few flies (<1%) emerged out of the *Spiroplasma*-uninfected treatments due to wasp attack, differences in fly survival across the three temperatures are explained by differences in temperature across the *Spiroplasma*-infected treatments only. Specifically, in the ‘pre-attack’ regime, *Spiroplasma*-infected fly larva-to-adult survival at 21°C was lower than at 23°C and 25°C (mean fly survival at 21°C: 24%, at 23°C: 46.3%, at 25°C: 56.7%, n = 10 vials in each case). Comparably, in the ‘constant’ temperature regime, fly larva-to-adult survival at 21°C was lower than at 23°C and 25°C (mean fly survival at 21°C: 20.3%, at 23°C: 39.3%, at 25°C : 57%, n= 10, 10 and 9 vials respectively). These data, including a lack of effect of temperature in the ‘attack and protection’ regime, indicate that cooler temperatures during maternal development and early adulthood lead to reduced strength of *Spiroplasma*-mediated fly protection in subsequent offspring under wasp attack.

**Table 2:** Analysis of deviance table for fly larva-to-adult survival for the ‘pre-attack’, ‘attack and protection’ and ‘constant’ temperature regimes in the wasp treatment.

| **Pre-attack** | Sum Sq | df | *F* | *p*-value |
| --- | --- | --- | --- | --- |
| Temperature | 69.31 | 2 | 33.67 | <0.001 |
| *Spiroplasma* infection | 625.87 | 1 | 608.04 | <0.001 |
| Temperature * *Spiroplasma* infection | 1.28 | 2 | 0.62 | 0.540 |
| Residuals | 54.55 | 53 |  |  |
| **Attack and protection** |  |  |  |  |
| Temperature | 6.32 | 2 | 1.67 | 0.199 |
| *Spiroplasma* infection | 1168.01 | 1 | 615.79 | <0.001 |
| Temperature * *Spiroplasma* infection | 1.32 | 2 | 0.35 | 0.708 |
| Residuals | 98.63 | 52 |  |  |
| **Constant** |  |  |  |  |
| Temperature | 87.94 | 2 | 25.87 | <0.001 |
| *Spiroplasma* infection | 559.37 | 1 | 329.14 | <0.001 |
| Temperature * *Spiroplasma* infection | 2.58 | 2 | 0.76 | 0.473 |
| Residuals | 91.77 | 54 |  |  |

*Note: A generalised linear model with logit link and quasibinomial fit was applied except for the Pre-attack (a) analysis in which a binomial fit was applied. (Attack and protection): The dispersion parameter is 1.90. (Constant): The dispersion parameter is 1.70.*

As expected, *Spiroplasma* infection had a significant positive effect on the fly larva-to-adult survival in the presence of wasps across all temperature regimes (Table 2). *Spiroplasma*-infected flies had higher survival than *Spiroplasma*-uninfected flies. However, the interaction between temperature and *Spiroplasma* infection was not found to be significant in any of the temperature regimes due to *Spiroplasma* consistently increasing fly survival across all temperatures (Figure 3A; B; C).

**Figure 3:**
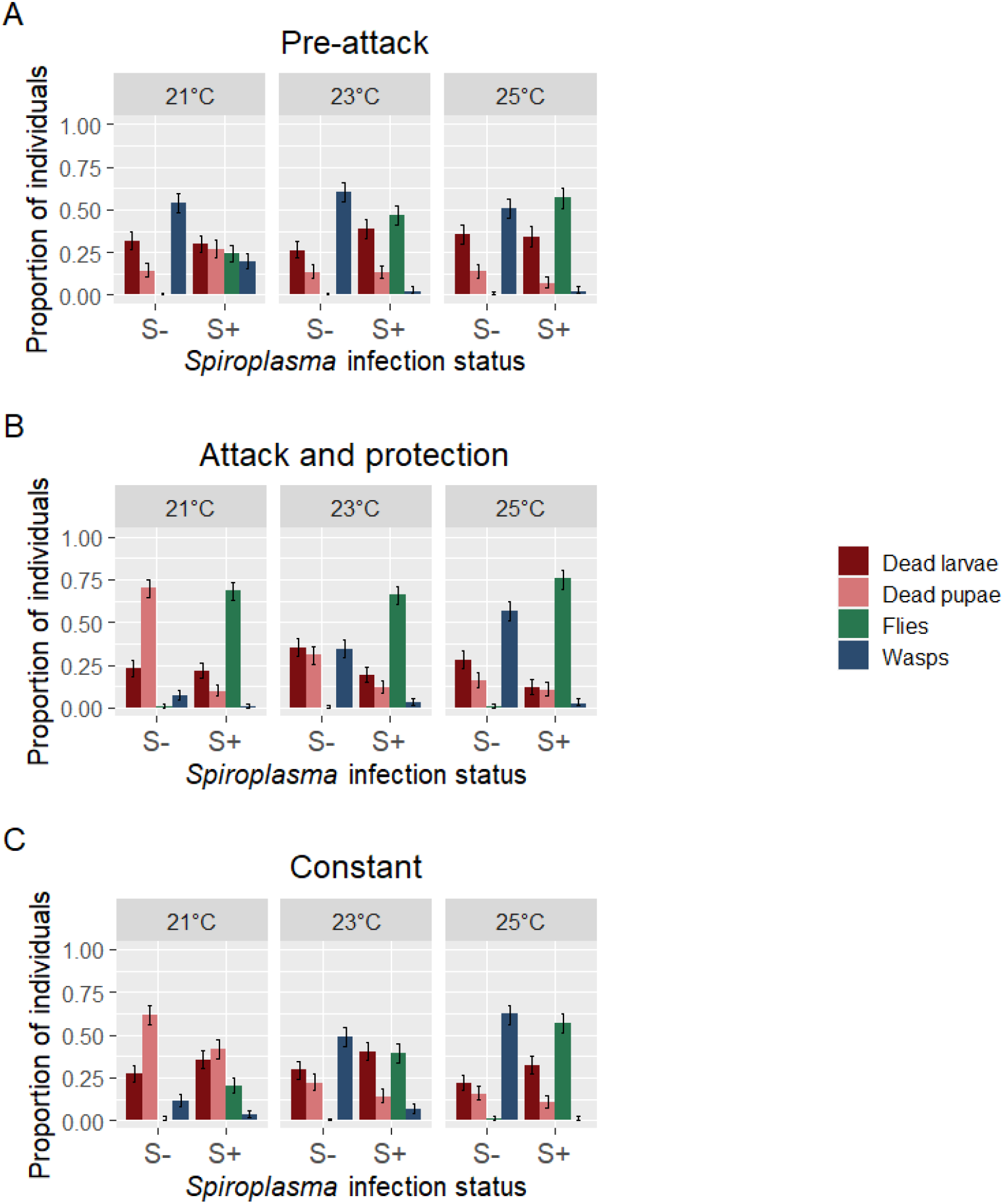
Proportion of dead larvae (red), dead pupae (pink), emerging flies (green) and emerging wasps (blue) for *Spiroplasma*-uninfected (S-) and -infected (S+) F1 *Drosophila melanogaster* under the wasp-attacked ‘pre-attack’ (A), ‘attack and protection’ (B) and ‘constant’ (C) temperature regimes. Error bars represent 95% binomial confidence intervals.

### Interaction between thermal environment, fly Spiroplasma infection and wasp mortality

Wasp mortality was also affected by temperature (Figure 3A; B; C). As very few wasps emerged out of the *Spiroplasma*-infected treatments (<7%), differences in wasp mortality across the three temperatures are largely driven by differences in temperature across the *Spiroplasma*-uninfected treatments only. In the ‘attack and protection’ temperature regime, wasp mortality in *Spiroplasma*-uninfected flies at 21°C was higher than at 23°C and 25°C (mean wasp mortality at 21°C: 93%, at 23°C: 66%, at 25°C: 44%, n = 10 vials in each case). Comparably, in the constant temperature regime, wasp mortality in *Spiroplasma*-uninfected flies at 21°C was higher than at 23°C and 25°C (mean wasp mortality at 21°C: 89%, at 23°C: 77%, at 25°C: 38%, n = 10 vials in each case). These data indicate that wasp mortality in *Spiroplasma*-uninfected flies is higher at cooler temperatures.

Temperature also had a significant effect on wasp mortality in the ‘pre-attack’ temperature regime (Table 3). This may appear surprising given that at this stage, only the flies had experienced experimental temperatures, whereas wasps had only experienced a temperature of 25°C across all treatments of the ‘pre-attack’ regime. This effect was mainly brought about by the high number of wasps emerging out of the 21°C *Spiroplasma*-infected treatment (mean wasp mortality = 80%) compared to 23°C (mean wasp mortality = 98%) and 25°C (mean wasp mortality = 98%) indicating potential segregation or low titre of *Spiroplasma* in flies at 21°C, resulting in lower wasp mortality (Figure 3B). However, this effect was not mirrored in the 21°C ‘constant’ regime, indicating that even in *Spiroplasma*-uninfected flies, wasp mortality is increased at cooler temperatures.

**Table 3:** Analysis of deviance table for wasp mortality for the ‘pre-attack’, ‘attack and protection’ and ‘constant’ temperature regimes in the wasp treatment.

| **Pre-attack** | Sum Sq | df | *F* | *p*-value |
| --- | --- | --- | --- | --- |
| Temperature | 19.24 | 2 | 3.83 | 0.028 |
| *Spiroplasma* infection | 492.59 | 1 | 196.02 | <0.001 |
| Temperature * *Spiroplasma* infection | 61.88 | 2 | 12.31 | <0.001 |
| Residuals | 133.19 | 53 |  |  |
| **Attack and protection** |  |  |  |  |
| Temperature | 186.83 | 2 | 40.79 | <0.001 |
| *Spiroplasma* infection | 342.72 | 1 | 149.64 | <0.001 |
| Temperature * *Spiroplasma* infection | 4.28 | 2 | 0.94 | 0.399 |
| Residuals | 119.10 | 52 |  |  |
| **Constant** |  |  |  |  |
| Temperature | 166.94 | 2 | 34.09 | <0.001 |
| *Spiroplasma* infection | 449.97 | 1 | 183.75 | <0.001 |
| Temperature * *Spiroplasma* infection | 46.81 | 2 | 9.56 | <0.001 |
| Residuals | 132.24 | 54 |  |  |

*Note: A generalised linear model with logit link and quasibinomial fit was applied. (Pre-attack): The dispersion parameter is 2.51. (Attack and protection): The dispersion parameter is 2.29. (Constant): The dispersion parameter is 2.45.*

As expected, wasp mortality was significantly affected by the *Spiroplasma* infection status of flies across all temperature regimes (Table 3). Wasp mortality was higher in *Spiroplasma*-infected flies compared to *Spiroplasma*-uninfected flies (Figure 3A; B; C). However, in the pre-attack temperature regime, the effect of fly *Spiroplasma* infection on wasp mortality was less at 21°C compared to 23°C or 25°C resulting in a significant interaction between *Spiroplasma* infection and temperature. Similarly, there was a significant interaction between fly *Spiroplasma* infection and temperature in the constant temperature regime. Here, the effect of *Spiroplasma* infection was greater at 25°C compared to 23°C or 21°C.

### The effect of temperature on sex ratio

The sex of each fly emerging from the *Spiroplasma*-infected 21°C, 23°C and 25°C treatments from each experimental temperature regime was recorded to determine the impact of temperature on the male-killing phenotype. In the ‘constant’ and ‘pre-attack’ temperature regimes, male survival increased from <2% at 25°C to 15-19% at 21°C (Table 4). However, in the presence of wasps, male survival was 0% for both 21°C and 25°C, indicating that wasps were killing all *Spiroplasma*-uninfected males produced (Table 4). The same effect was observed in the attack and protection treatment where sporadic males were produced in each temperature treatment in the absence of wasps, but were absent in wasp treatments, presumed to have been eliminated by wasps (Table 4). Consistent with this interpretation, a subset of male flies emerging out of the un-attacked constant temperature regime at 21°C (3 samples) and at 23°C (8 samples) were screened for *Spiroplasma* infection using methods described in (Jones and Hurst, 2021b) and all (11/11) were found to be uninfected.

**Table 4:** Sex ratio of *Spiroplasma*-infected *Drosophila melanogaster* emerging from the 21°C, 23°C and 25°C treatments in the ‘pre-attack’, ‘attack and protection’ and ‘constant’ temperature regimes.

| Temperature regime | Temperature | Wasp treatment | Total fly | Total female | Total male | % male |
| --- | --- | --- | --- | --- | --- | --- |
| Pre-attack | 25°C | Lb- | 238 | 235 | 3 | 1 |
|  |  | Lb+ | 153 | 153 | 0 | 0 |
|  | 23°C | Lb- | 205 | 196 | 9 | 4 |
|  |  | Lb+ | 139 | 139 | 0 | 0 |
|  | 21°C | Lb- | 218 | 177 | 41 | 19 |
|  |  | Lb+ | 72 | 72 | 0 | 0 |
| Attack and protection | 25°C | Lb- | 243 | 239 | 4 | 2 |
|  |  | Lb+ | 181 | 181 | 0 | 0 |
|  | 23°C | Lb- | 186 | 183 | 3 | 2 |
|  |  | Lb+ | 198 | 198 | 0 | 0 |
|  | 21°C | Lb- | 206 | 202 | 4 | 2 |
|  |  | Lb+ | 183 | 183 | 0 | 0 |
| Constant | 25°C | Lb- | 234 | 233 | 1 | 0 |
|  |  | Lb+ | 171 | 171 | 0 | 0 |
|  | 23°C | Lb- | 210 | 205 | 5 | 2 |
|  |  | Lb+ | 118 | 118 | 0 | 0 |
|  | 21°C | Lb- | 247 | 209 | 38 | 15 |
|  |  | Lb+ | 61 | 61 | 0 | 0 |

### The effect of temperature and wasp attack on *Spiroplasma* titre

As expected, *Spiroplasma*-infected flies reared at cooler temperatures were found to have lower *Spiroplasma* titre than flies reared at warmer temperatures (Figure 4). Flies reared at 21°C were found to have 2.05× lower *Spiroplasma* titre than flies reared at 23°C and 3.08× lower *Spiroplasma* titre than flies reared at 25°C (21°C mean = 0.81, n = 32; 23°C mean = 1.65, n = 39; 25°C mean = 2.48, n = 37;*χ^2^=* 27.012, d.f. = 2, *p* < 0.001). There was no significant overall effect of wasp attack on *Spiroplasma* titre (*χ*^2^= 0.169, d.f. = 1, *p* = 0.681), nor a significant interaction between temperature and wasp presence (*χ*^2^= 1.960, d.f. = 2, *p* = 0.375). *Spiroplasma*-negative flies, as determined by qPCR, were observed in the 21°C non-wasp attacked treatment but not in other treatments (21°C Lb-: 8/21 *Spiroplasma* negatives vs. 21°C Lb+: 0/19 *Spiroplasma* negatives), indicating that *Spiroplasma* infection segregates at 21°C, but uninfected individuals created were subsequently killed by wasps in the wasp-attack treatment. Individuals found to be uninfected were not included in this analysis.

**Figure 4:**
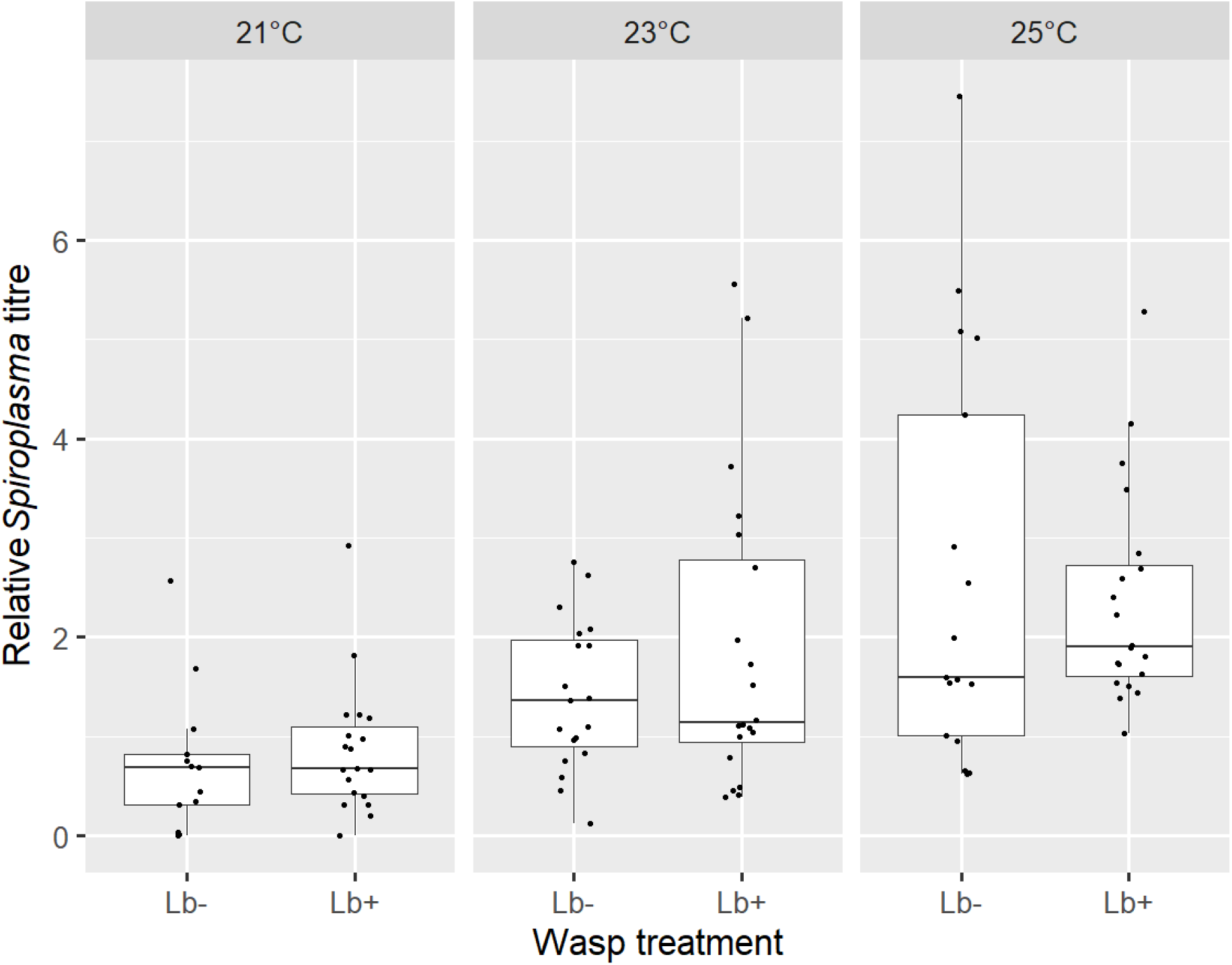
Relative *Spiroplasma* titre of wasp-attacked (Lb+) and no wasp attack control (Lb-) *Spiroplasma*-infected *Drosophila melanogaster* emerging from the 21°C, 23°C and 25°C treatments in the constant temperature regime. The box plots display the upper and lower quartiles, the median and the range. Each dot represents a single fly.

### The effect of temperature on wasp oviposition

The average number of wasp eggs laid into a fly larva across a 48 h period of parasitisation was >3 but <4 for all treatments (Figure 5). There was no significant effect of temperature on the number of wasp eggs laid into fly larvae (*χ^2^* = 2.52, d.f. = 2, *p =* 0.284).

**Figure 5:**
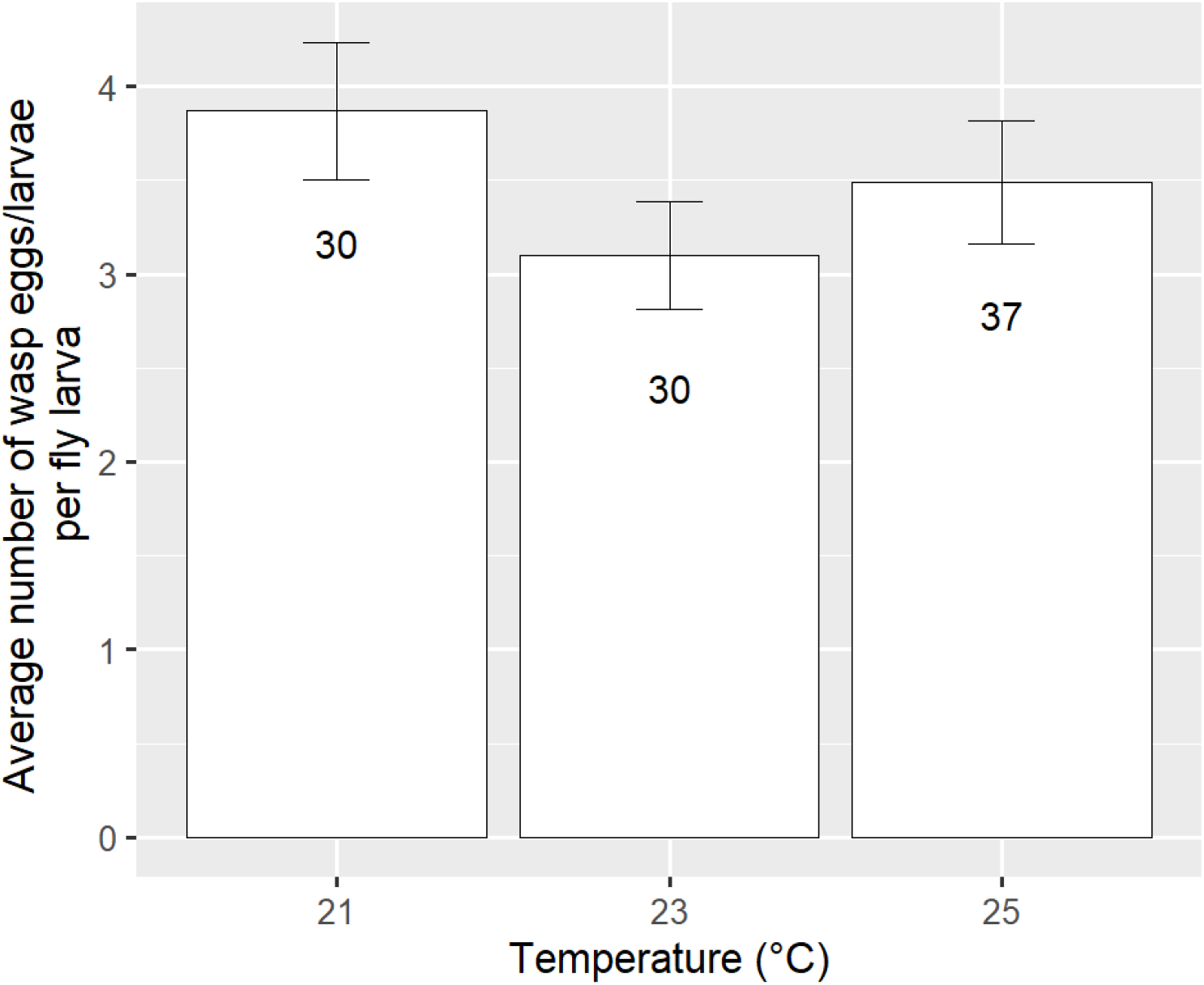
The average number of wasp eggs/larvae in *Drosophila melanogaster* (Canton-S) larvae following 48 h of parasitisation by *Leptopilina boulardi* (Lb17) at three different temperatures: 21°C, 23°C and 25°C. Error bars depict ± SE.

### Further analysis on the effect of temperature on fly survival and wasp mortality

It is apparent from the data above that *Spiroplasma* starts to segregate from flies at 21°C. To determine temperature effects on fly survival in cases where *Spiroplasma* is retained, we repeated the fly survival analysis using only data from the 23°C and 25°C treatments where *Spiroplasma* transmission is still highly efficient. In line with the previous analysis (Table 2), temperature was found to have a significant effect on the fly larva-to-adult survival in the presence of wasps, in both the pre-attack and constant temperature regimes but not in the post-attack temperature regime (Table S2). This result indicates that low *Spiroplasma* titre can reduce fly larva-to-adult survival under wasp attack.

Whilst it is not possible to precisely partition the impact of *Spiroplasma* loss and low titre on the reduced protection phenotype seen when flies are exposed to 21°C at the pre-attack stage, it appears likely that the effect derives from a combination of *Spiroplasma* loss in flies and *Spiroplasma* infected flies surviving attack at a lower rate. Within the ‘pre-attack’ treatment, 41 males and 177 females emerged out of the no wasp control treatment (Table 4). As these males were found to be uninfected, we can assume that a similar number of females were also uninfected and had likely lost infection at the point of embryogenesis (41 males + 41 females = 82 flies uninfected out of a total of 218 total flies; 82/218 = c. 37% uninfected). Assuming fly survival can only occur in the presence of *Spiroplasma*, uninfected larvae should be discounted, such that the observed fly survival of 24% in the 21°C pre-attack *Spiroplasma*-infected treatment (Figure 3A) should be normalised to the 63% flies that are infected (100% - 37% uninfected flies). Thus, infected flies that have experienced 21°C prior to attack have a c. 38% chance of surviving attack in the presence of *Spiroplasma*, compared to fly survival of 46% and 57% at 23°C and 25°C respectively during the pre-attack period.

For wasp mortality, fly controls without *Spiroplasma* in the 21°C pre-attack regime showed a wasp mortality rate of 46% (Figure 3A). We know that 37% of flies in the *Spiroplasma*-infected attacked treatment are in fact uninfected, and in these we expect 46% wasp mortality, such that if *Spiroplasma* kills all wasps we expected wasp survival of around 20% (mortality around 80%).

This value parallels the actual 80% wasp mortality observed (20% survival) (Figure 3A), leading to the conclusion that wasps only complete development in *Spiroplasma*-uninfected individuals i.e. low titre *Spiroplasma* is sufficient to kill wasps, even if it is not sufficient to rescue the fly host. It may be that a low density of *Spiroplasma* is enough to kill developing wasps, but this takes longer, allowing the wasp to cause more damage to the fly resulting in reduced fly rescue.

## Discussion

The outcome of natural enemy attack in insects can be modulated by the presence of defensive heritable symbionts residing within the host. Beyond symbiont presence, studies have shown that genetic and environmental factors can also mediate the outcome of symbiont-mediated defence (Doremus *et al*., 2018; Vorburger and Perlman, 2018; Corbin *et al*., 2021; Chrostek *et al*., 2021). Thermal environment is of particular importance to symbiont-mediated traits, with minor differences in temperature driving major changes in the outcome of the interaction (Higashi *et al*., 2020). In this study we examined the thermal sensitivity of protection against wasp attack. Following recent studies of *Wolbachia*, we determined the importance of current vs historical thermal environment and concluded that historical environment impacted the protection phenotype in the *D. melanogaster-Spiroplasma-L. boulardi* interaction, but the thermal environment during attack/defence itself was unimportant.

Only the temperature before wasp attack was observed to impact *Spiroplasma*-mediated protection, measured in terms of fly survival. This inference derives from reduction in fly survival in the pre-attack lower temperatures treatments, the comparability of effect size of the ‘pre-attack’ regime with that seen in the ‘constant’ regime, and the absence of any thermal effect on protection when lower temperatures were experienced solely during attack/defence phases (attack and protection temperature regime). The absence of any effect of temperature on *Spiroplasma*-mediated fly survival during or after wasp attack suggests that thermal effects observed are mediated predominantly by historical thermal effects on *Spiroplasma* titre and not by thermally dependent properties of the wasp or *Spiroplasma* defence molecules themselves. In support of this, relative *Spiroplasma* titre of F1 adults, reared through 21°C in the constant temperature regime, was found to be ~68% lower than in adults reared through the 25°C treatment. A second impact of temperature was on the presence of *Spiroplasma* at lower temperatures: evidence from qPCR indicates that *Spiroplasma* infection was segregating at 21°C. Collectively, these results demonstrate that lower *Spiroplasma* titre and loss of infection are predominately responsible for the weak *Spiroplasma*-mediated protection observed at 21°C. This result is perhaps unsurprising given the repeated evidence that *Spiroplasma* is a cool sensitive symbiont, whereby its transmission efficiency and phenotype expression are negatively impacted by lower temperatures (Montenegro and Klaczko 2004; Corbin *et al*., 2021; Osaka *et al*., 2008). Nevertheless, it remains possible that temperature may be additionally affecting other properties of *Spiroplasma* defence, as is suspected in cool sensitivity of *Wolbachia* protection against *Drosophila* C virus (Chrostek *et al*., 2021).

As the pre-attack treatment was virtually a single generation and only included a single day (egg development) of the second generation, it is most parsimonious to conclude the effect of temperature on *Spiroplasma*-mediated defence is transgenerational. The thermal environment experienced by the mother determines the outcome of *Spiroplasma*-mediated protection in her daughters. This effect is likely mediated by mothers with reduced *Spiroplasma* titre transmitting lower *Spiroplasma* densities to their offspring, subsequently leading to reduced protection from wasps. Transgenerational influences of this kind have shown in the *Drosophila innubila-male*-killing *Wolbachia* symbiosis where female *Drosophila innubila* infected with low *Wolbachia* titres produce daughters with low *Wolbachia* titre that themselves exhibit weaker male-killing amongst their progeny (Dyer and Jaenike, 2005).

*Spiroplasma* in *D. melanogaster* has only been recorded in the tropics and has a much narrower geographical range than the comparable defensive symbiosis in *D. hydei*, in which the symbiont is also recorded in temperate regions (Montenegro *et al*., 2005; Pool *et al*., 2006; Kageyama *et al*., 2006). Our data – indicating segregation and loss of *Spiroplasma* at 21°C and reduction in protection at 23°C in *D. melanogaster*– correlate with this narrow range; contrastingly the infection in *D. hydei* (present in temperate areas) segregates only after a number of generations at 15°C and protection is ablated at 18°C. An emerging question is what determines the variation we see in the thermal sensitivity of *Spiroplasma*-insect symbiosis, and to what extent is thermal sensitivity evolvable?

In summary, our work has extended our understanding of how the thermal environment affects *Spiroplasma*-mediated protection in *Drosophila*. Our results reveal that the developmental thermal environment of the mother is more important to the outcome of *Spiroplasma*-mediated protection against wasps than the thermal environment of individuals during or after attack. This finding has implications for *Spiroplasma* dynamics in natural populations. Not only should *Spiroplasma* dynamics be considered in relation to the strain of circulating parasitoid wasp, but also the historical thermal environment the flies have been exposed to. Indeed, it should be noted that our study was only conducted on one strain of *L. boulardi* wasp and that more complex interactions between the thermal environment and wasp strain/species may exist (G x G x E interactions). Future work should consider the effect of the thermal environment on other strains of *Leptopilina* wasp to understand whether more complex G x E interactions are important in this system.

## Acknowledgments

We would like to thank Dr Todd Schlenke for providing the Lb17 wasp strain. We would also like to thank Helen Davison for her help picking larvae for the experiments. We thank Dr Ewa Chrostek, Prof Andrea Betancourt and Prof Nina Weddell for their kind comments on the manuscript. This project was supported by funding from the NERC (Studentship to J.J., grant number NE/L002450/1) and a NERC grant award (NE/V011979/1).

## Data accessibility

Data generated and analysed during this study are available at figshare (https://doi.org/10.6084/m9.figshare.c.6176527.v1).

## Appendix

**Table S1:**
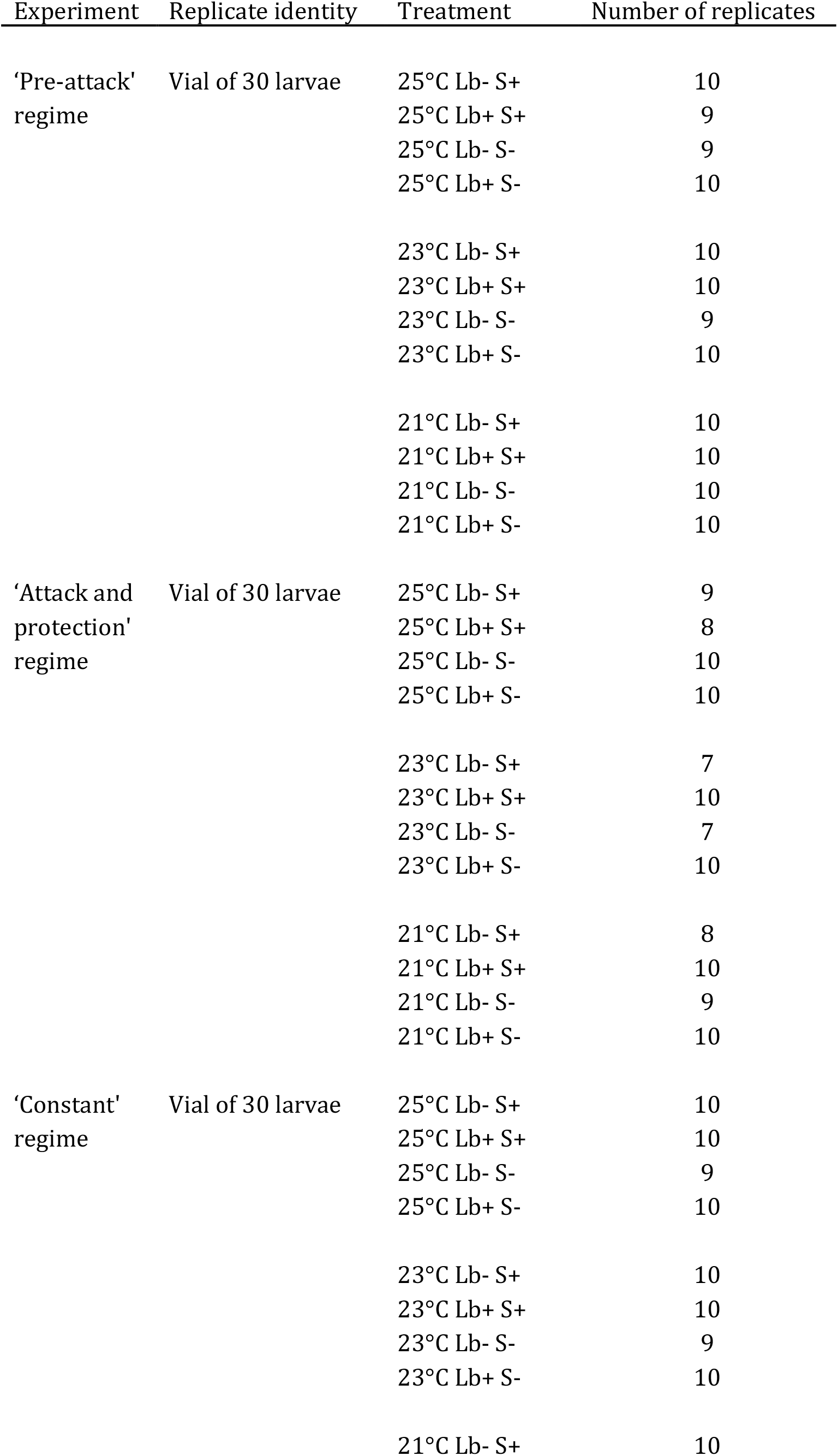
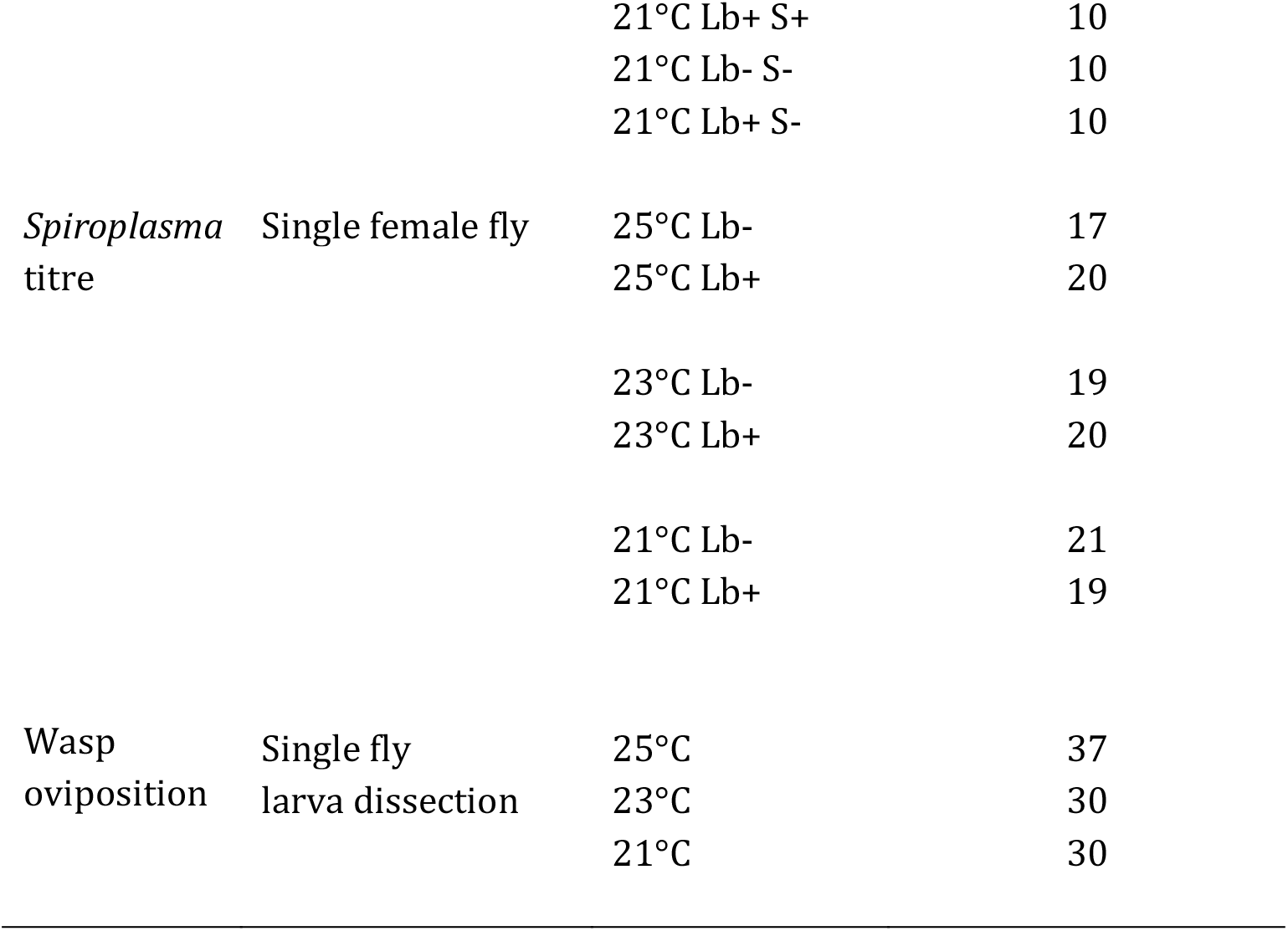
Replicate identity and number for chapter 4 experiments (Lb- and Lb+: wasp presence and absence; S- and S+: *Spiroplasma* presence and absence).

**Table S2:** Analysis of deviance table for fly larva-to-adult survival for the ‘pre-attack’, ‘attack and protection’ and ‘constant’ temperature regimes in the wasp treatment at 23°C and 25°C only.

| **Pre-attack** | Sum Sq | df | F | p-value |
| --- | --- | --- | --- | --- |
| Temperature | 6.46 | 1 | 5.27 | 0.028 |
| *Spiroplasma* infection | 516.48 | 1 | 421.06 | <0.001 |
| Temperature * *Spiroplasma* infection | 1.01 | 1 | 0.82 | 0.370 |
| Residuals | 42.93 | 35 |  |  |
| **Attack and protection** |  |  |  |  |
| Temperature | 6.12 | 1 | 2.73 | 0.108 |
| *Spiroplasma* infection | 784.49 | 1 | 349.68 | <0.001 |
| Temperature * *Spiroplasma* infection | 0.98 | 1 | 0.44 | 0.514 |
| Residuals | 76.28 | 34 |  |  |
| **Constant** |  |  |  |  |
| Temperature | 477.94 | 1 | 10.10 | 0.003 |
| *Spiroplasma* infection | 20.04 | 1 | 240.83 | <0.001 |
| Temperature * *Spiroplasma* infection | 1.59 | 1 | 0.80 | 0.377 |
| Residuals | 71.45 | 36 |  |  |

Note: A generalised linear model with logit link and quasibinomial fit was applied except for the Pre-attack analysis in which a binomial fit was applied. (attack and protection); The dispersion parameter is 2.24. (constant); The dispersion parameter is 1.98.

